# Compensatory reconfiguration of functional networks between white matter and grey matter in Alzheimer’s disease

**DOI:** 10.1101/2021.08.30.458154

**Authors:** Xingxing Zhang, Qing Guan, Debo Dong, Fuyong Chen, Jing Yi, Yuejia Luo, Haobo Zhang, Alzheimer’s Disease Neuroimaging Initiative

**Author notes:** **Corresponding author** Haobo Zhang, Center for Brain Disorders and Cognitive Sciences, Shenzhen University, Shenzhen, China, mailing address: No. 3688 Nanhai Ave., Nanshan District, Shenzhen 518060, China. **Declarations of interest: none**.

## Abstract

The temporal synchronization of BOLD signals within white matter (WM) and between WM and grey matter (GM) exhibited intrinsic architecture and cognitive relevance. However, few studies examined the network property within- and between-tissue in Alzheimer’s disease (AD). The hub regions with high weighted degree (WD) were prone to the neuropathological damage of AD. To systematically investigate the changes of hubs within- and between-tissue functional networks in AD patients, we used the resting-state fMRI data of 30 AD patients and 37 normal older adults (NC) from the ADNI open database, and obtained four types of voxel-based WD metrics and four types of distant-dependent WD metrics (ddWD) based on a series of Euclidean distance ranges with a 20mm increment. We found that AD patients showed decreased within-tissue ddWD in the thalamic nucleus and increased between-tissue ddWD in the occipito-temporal cortex, posterior thalamic radiation, and sagittal stratum, compared to NC. We also found that AD patients showed the increased between-tissue FCs between the posterior thalamic radiation and occipito-temporal cortex, and between the sagittal stratum and the salience and executive networks. The dichotomy of decreased and increased ddWD metrics and their locations were consistent with previous studies on the neurodegnerative and compensatory mechanisms of AD, indicating that despite the disruptions, the brain still strived to compensate for the neural inefficiency by reorganizing functional circuits. Our findings also suggested the short-to-medium ranged ddWD metrics between WM and GM as useful biomarker to detect the compensatory changes of functional networks in AD.

## 1. Introduction

Neural activity proceeds with the participation of neurons and fiber. In this process, the information transfers among different neurons via the dendrites and axons. However, the neural activities in the white matter (WM) were often ignored, possibly due to its relatively weak signal than grey matter (GM) and lack of understanding of its function (Gawryluk, Mazerolle, & D’Arcy, 2014). It was suggested that the blood oxygen level-dependent (BOLD) signal in the WM were originated from neurons, which could bring a series of biochemical changes, e.g. the exchanges of potassium and calcium ions and the production of nitric oxide. These changes might lead to vascular changes in the WM and increased BOLD signal (Gawryluk et al., 2014; Marcar & Loenneker, 2004; Smith et al., 2002). Previous studies also showed that the BOLD fluctuations detected in the WM were responsive to the task stimuli (Ding et al., 2018; Huang et al., 2018; Huang et al., 2020).

The temporal synchronization of WM BOLD signals in the task-related WM regions was enhanced during performing various cognitive tasks (Huang et al., 2020; Marussich, Lu, Wen, & Liu, 2017). At rest, the functional network within the WM demonstrated an intrinsic architecture, consistent with the structural organization of WM tracts (Huang et al., 2018; Peer, Nitzan, Bick, Levin, & Arzyt, 2017). Recently, the functional network of WM has received much attention for its applications in schizophrenia, attention-deficit/hyperactivity disorder, and mild cognitive impairment (Bu et al., 2020; X. Chen et al., 2017; Jiang et al., 2019).

A few studies examined the temporal synchronization of BOLD signals between WM bundles and GM regions in human and non-human primates, and found that the spatial distribution pattern of functional connectivity (FC) between them was not random, but in a similar arrangement as the anatomical construction that they partook in (Ding et al., 2018; Huang et al., 2020; Marussich et al., 2017; Peer et al., 2017; Wu et al., 2019). The FC strength between the two tissues was increased with the cognitive load in different tasks (Ding et al., 2018; Huang et al., 2020; Marussich et al., 2017).

As the temporal synchronization of WM has become a promising neuroimaging biomarker, two recent studies used the FCs between 48 WM tracts and 82 regional GM volumes to classify Alzheimer’s disease (AD) from normal older adults (Gao et al., 2020; Zhao, Ding, Du, Wang, & Men, 2019). However, the topological changes in the functional network of WM in AD remained unclear. Previous studies showed that the hubs of functional network in the GM were specifically susceptible to the neurodegenerative damage of AD (Buckner et al., 2009; de Haan, Mott, van Straaten, Scheltens, & Stam, 2012). Possibly, the hubs of the WM functional networks also changed in AD patients.

From abundant functional network metrics, we chose weighted degree (WD) as the hub regions with high WD were especially prone to AD damage. WD, similar to the metrics of functional connectivity strength or density used in previous studies (Z. J. Dai et al., 2015; Tomasi & Volkow, 2010), was an indicator of the total strength of the FCs of a node with other nodes (Rubinov & Sporns, 2010; Zuo et al., 2012). Voxel-wise parcellation allows the neural signals more homogeneous extracted from the fine-grained units, the network more prone to the small-world property, and the results in more precise locations (de Reus & van den Heuvel, 2013; Hayasaka & Laurienti, 2010). According to the locations of two nodes for a FC, there would be four types of voxel-based WDs: WD-G as the weighted sum of FCs of a given GM voxel with other GM voxels, WD-W of a given WM voxel with other WM voxels, WD-GW of a given GM voxel with WM voxels, and vice versa for WD-WG. Moreover, previous studies showed that the FC was anatomical distance-dependent, with the short-ranged and long-ranged FCs differing on the metabolic costs and the susceptibility to neurodegenerative diseases (Z. J. Dai et al., 2015; Liang, Zou, He, & Yang, 2013). Hence, we also obtained the distant-dependent WD metrics (ddWD-W, ddWD-G, ddWD-GW, and ddWD-WG) according to the Euclidean distance between two voxels. Through a series of group comparison analysis on the four types of voxel-based WD and ddWD metrics between AD and normal controls (NC), we aimed to investigate the changes of WM functional networks in AD patients and hypothesized that the changes might be consistent with the neuropathological mechanism of AD.

## 2. Materials and Methods

### 2.1 ADNI database

The present study used the open database, the Alzheimer’s Disease Neuroimaging Initiative (ADNI) to acquire data. ADNI was launched in 2004 to investigate AD and its prodromal stages (*http://www.adniinfo.org*). ADNI included four databases (ADNI-1, ADNI-2, ADNI-3, ADNI-GO). An initial five-year study termed ADNI-1, was followed by two renewal studies termed ADNI-2 and -3 (Weiner et al., 2017). As the ADNI-1 phase did not collect resting-state fMRI data, we acquired data from the ADNI-2. According to the standardized protocol, the ADNI data was collected from various acquisition sites across Canada, and the United States, approved by the Institutional Review Board at each acquisition site, and written informed consent was obtained from each participant.

### 2.2 Participants

The participants with the following criteria were included in this study: (1) a clinician-confirmed diagnosis of AD or ‘‘normal’’ at the screening visit; and (2) the complete resting-state fMRI data available for participants at their first scan time in the ADNI 2 database. In total, 30 AD patients and 37 NC were included in this study. No significant differences were found between the two groups on age and gender

### 2.3 Image acquisition and preprocessing

All participants were scanned with a 3.0 T Philips scanner (Jack et al., 2008). The resting-state fMRI data were obtained using an echo-planar imaging (EPI) sequence with the following parameters: repetition time (TR) = 3000 ms, echo time (TE) = 30 ms, flip angle = 80°, number of slices = 48, slice thickness = 3.313 mm, voxel size = 3 mm × 3 mm × 3 mm, voxel matrix = 64 × 64, and total volume = 140.

Rs-fMRI images were preprocessed using the Data Processing and Analysis of Brain Imaging software package (DPABI, *http://rfmri.org/dpabi*) and the SPM12. The preprocessing procedure consisted of eight steps: 1) discarding the first ten volumes, 2) slice-timing, 3) head motion correction (the maximum motion threshold ≤2mm or 2°), 4) normalization to the MNI template and resampled to 3 × 3 × 3 mm^3^, 5) detrend, 6) nuisance covariates regression (Friston 24 for head motion, global signal, and cerebrospinal fluid signal regression), 7) temporal scrubbing (the scan volume with the frame-wise displacement (FD) >1 was removed), and 8) band-pass filter (0.01 Hz– 0.15 Hz) to reduce low-frequency drift and high-frequency physiological noise (Jiang et al., 2019; Peer et al., 2017). One AD participant was removed from this study due to the maximum head motion beyond the threshold.

### 2.4 Network construction and distance-dependent weighted degree (ddWD)

#### 2.4.1 Functional network in GM (ddWD-G)

The calculation pipeline of four ddWD metrics were demonstrated in Fig 1. The GM mask was generated from the GM probability map (provided by SPM12) for the voxels with a GM probability value >0.2 (Du et al., 2015). The cut-off value on Euclidean distance between two voxels was often set arbitrarily as 75mm (Deng et al., 2016; Sheng et al., 2018) or incrementally with 10mm (Z. J. Dai et al., 2015). In this study, the incremental unit of 20mm was used to divide the distance of voxels in the GM mask. Hence, there were nine ranges of Euclidean distance between any two GM voxels, starting from 0 to 180mm with a 20mm increase for each range. According to the procedure of calculating WD in previous studies (Liang et al., 2013; H. B. Zhang et al., 2018), the ddWD-G at a certain range was calculated as the sum of Fisher z-transformed Pearson’s correlation coefficients (Rs, the threshold for effective correlation>0.2) of any given voxel in the GM mask with all other GM voxels whose Euclidean distance to this voxel was within that range (Buckner et al., 2009; Z. J. Dai et al., 2015; H. B. Zhang et al., 2018). The voxel-based ddWD-G metrics at the nine ranges were smoothed (4mm) and then z-standardized.

**Fig 1.**
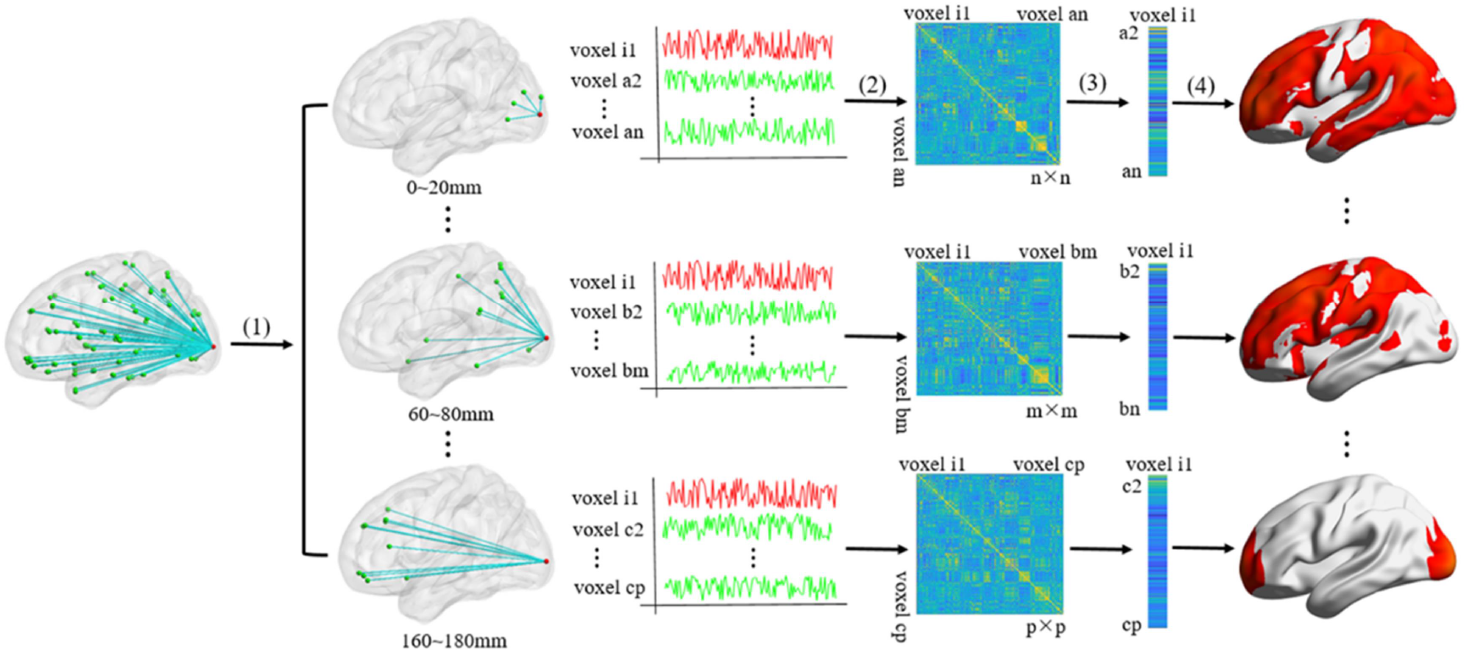
The calculation pipeline of ddWD. (1) The Euclidean distance between any voxel (e.g. Voxel i) and all other voxels in a relevant mask were calculated based on three-dimensional coordinates of these voxels. Then, the incremental unit of 20mm was used to divide the distance between voxels. (2) For the ddWD at a certain range, the Pearson’s correlation coefficients (Rs) of Voxel i with all other voxels whose Euclidean distance to this voxel was within that range were calculated, and then Fisher z-transformed (Rs in the diagonal elements and Rs ≤0.2 were set to 0). (3) The transformed Rs in a column for Voxel i were summed and referred as the ddWD for that voxel. (4) Steps (1) to (3) were repeated for all voxels in the relevant mask to obtain voxel-based ddWD. The voxel-based ddWD map at each Euclidean distance range was smoothed (4mm) and then z-standardized.

#### 2.4.2 Functional network in WM (ddWD-W)

The WM mask was generated from the WM probability map (provided by SPM12) for the voxels with a WM probability value >0.6 (Jiang et al., 2019; Peer et al., 2017). We excluded the voxels (n=4405) from the calculation of ddWD-G and ddWD-W, as these voxels were overlapped on the GM and WM masks, with a GM probability >0.2 and WM probability >0.6. These voxels were located in the boundary between GM and WM, as shown in Supplementary Fig s1. We used eight ranges to define Euclidean distance between any two voxels in the WM mask, starting from 0 to 180mm with a 20mm increase for each range. The ddWD-W at a certain range was the sum of Fisher z-transformed Rs of any given voxel in the WM mask with all other WM voxels whose Euclidean distance to this voxel was within that range. The ddWD-W metrics at eight ranges were smoothed (4mm) and then z-standardized.

#### 2.4.3 Functional networks between WM and GM (ddWD-GW and ddWD-WG)

We used nine ranges to define Euclidean distance between a voxel in the GM mask and a voxel in the WM mask, starting from 0 to 180mm with a 20mm increase for each range. The ddWD-GW at a certain range was the sum of the Fisher z-transformed Rs of any given voxel in the GM mask with all the voxels in the WM mask whose Euclidean distance to that GM voxel was within that range. The ddWD-GW metrics at nine ranges were smoothed (4mm) and then z-standardized.

The ddWD-WG at a certain range was the sum of the Fisher z-transformed Rs of any given voxel in the WM mask with all the voxels in the GM mask whose Euclidean distance to that WM voxel was within that range. The ddWD-WG metrics at nine ranges were smoothed (4mm) and then z-standardized.

#### 2.4.4 Weighted degree (WD) and mean maps

Before calculating ddWD metrics according to different ranges of Euclidean distance, we generated four types of WD metrics (WD-G, WD-W, WD-GW, and WD-WG) without dividing the distance between two voxels. In brief, the Rs of any given voxel with other voxels in relevant masks were calculated, Fisher r-to-z transformed, and then summed. The four voxel-based WD metrics for each individual were then smoothed (4mm) and z-standardized. To demonstrate the spatial distribution of the voxels with relatively high WD values (i.e. z-standardized value > 0.01), we calculated the mean voxel-based WD for the AD group and the NC group, respectively, and superimposed the mean maps of each group on 3D brain template (Fig 2). Similarly, the mean voxel-based ddWD metrics at each range for each group were calculated, and superimposed on 3D brain template (Fig 3).

**Fig 2.**
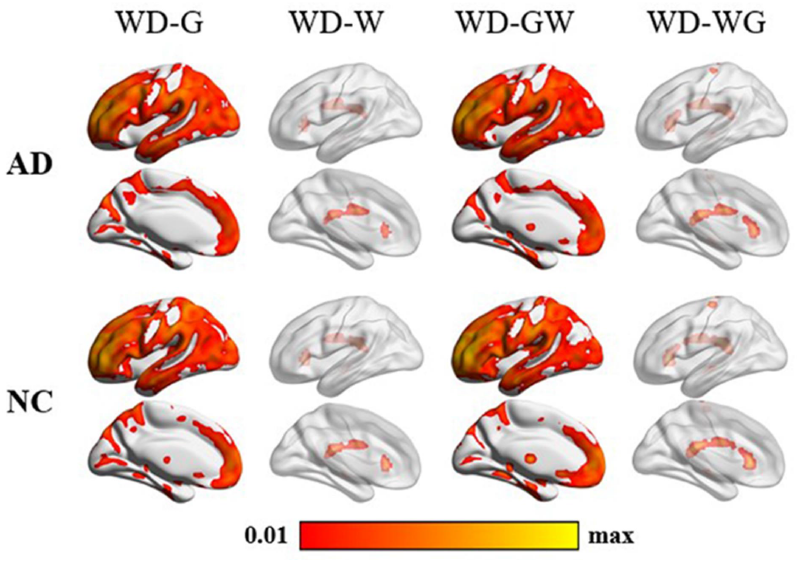
Mean maps of voxel-based weighted degree in AD and normal controls. The spatial distribution of hubs that were the voxels with high WD value, i.e. Z-score ranging 0.01∼maximum (note: the maximum value varied for each WD metric) were demonstrated. We calculated the mean voxel-based WD for the AD group and the NC group, respectively, and superimposed the mean maps on 3D brain template. In each group, four mean maps for four WD metrics were generated, including WD-G, WD-W, WD-GW, and WD-WG. The color bar represents the mean value, with higher values in warmer colors.

**Fig 3.**
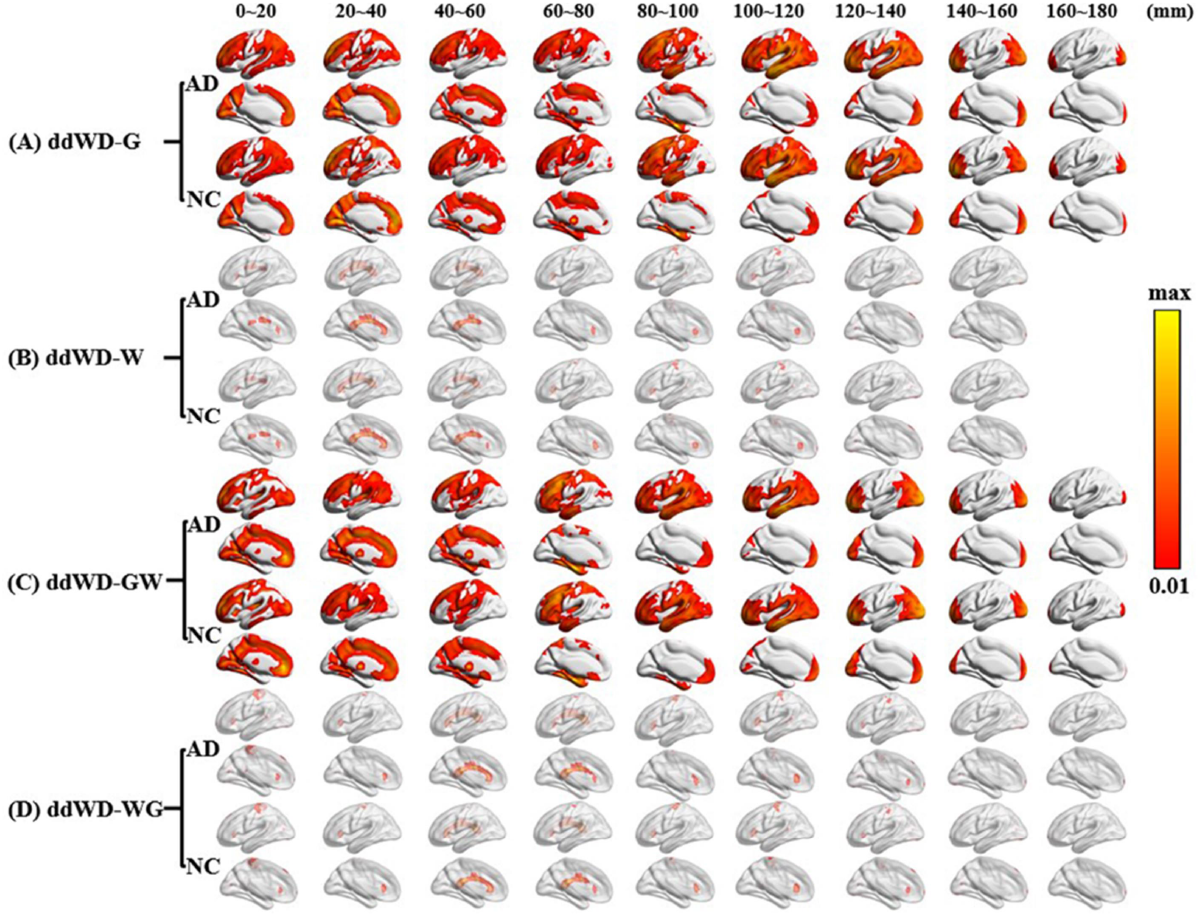
Mean maps of voxel-based distance-dependent weighted degree in AD and normal controls. The spatial distribution of hubs that were the voxels with high ddWD value, i.e. Z-score ranging 0.01∼maximum (note: the maximum value varied for each ddWD metric) were demonstrated. We calculated the mean voxel-based ddWD for the AD group and the NC group, respectively, and superimposed the mean maps on 3D brain template. A) the mean maps of ddWD-G metrics; B) the mean maps of ddWD-W; C) the mean maps of ddWD-GW metrics; and D) the mean maps of ddWD-WG metrics. The color bar represents the mean value, with higher values in warmer colors.

### 2.5 Seed-based functional connectivity

Compared various ddWD metrics of AD patients with NC would generate the supra-threshold clusters, where the ddWDs of a cluster of voxels were significantly smaller (or larger) in AD than NC. Using the supra-threshold cluster as the seed, we attempted to pinpoint the specific voxels whose FCs with the seed were also significantly smaller (or larger) in AD than NC.

Using the pre-processed rs-fMRI images, we firstly extracted voxel-wise time series from each seed and then calculated the average time series with the DPABI software (Data Processing & Analysis for Brain Imaging, *http://rfmri.org/dpabi*) (Chao-Gan & Yu-Feng, 2010). Secondly, the average time series of the seed was correlated with the time series of other voxels within either the GM or WM mask, dependent on the metrics of the seeds, i.e., GM mask for ddWD-G, WM mask for ddWD-W, WM mask for ddWD-GW, GM mask for ddWD-WG. Thirdly, as the seed was obtained from a ddWD metric at a certain Euclidean distance range, we only retained the FCs of the voxels whose Euclidean distance to any voxel of the seed was within that range. The resultant seed-based FCs then underwent the Fisher r-to-z transformation and smoothed (4mm).

### 2.6 Statistical analysis

We compared the differences between AD and NC on four types of WD (WD-G, WD-W, WD-GW, and WD-WG) and ddWD metrics (ddWD-G, ddWD-W, ddWD-GW, and ddWD-WG), implemented in SPM12. Using the resultant supra-threshold clusters as the seeds, we obtained the seed-based FCs and compared them between AD and NC. The controlled covariates for these comparison analyses were age, gender, and mean FD of head motion. The significance threshold was set at a voxel-level threshold of p<0.005 (uncorrected) combined with a cluster-level threshold of p<0.05 (FWE-corrected), based on the Gaussian Random Fields theory to correct for multiple comparisons.

## 3. Results

We did not find any significant difference between AD and NC on voxel-based WD metrics. However, the analysis with voxel-based ddWD showed significant results. Additionally, to check if the GM atrophy of AD patients was located in the classic hippocampal region, we compared the voxel-based GM volumes between the two groups using the T1-weighted images, and showed the method and results in the supplementary materials (Table s1, Fig s2).

### 3.1 Voxel-based ddWD-G

Compared to NC, AD patients showed smaller ddWD-G at 0∼20mm in the bilateral thalamus. Using this cluster as the seed, we did not find any significant difference between the two groups on the seed-based FCs (Table 2, Fig 4).

**Table 1.** Demographic characteristics and neuropsychological test results.

| Characteristic | NC (n=37) | AD (n=30) | Test statistic | P-value |
| --- | --- | --- | --- | --- |
| Age (year) | 75.84±7.09 | 73.14±6.51 | T= -1.59 | 0.117<sup>a</sup> |
| Sex (male/female) | 13/16 | 14/23 | $\chi^2=0.329$ | 0.566<sup>b</sup> |
| MMSE (NC/AD n=31/24) | 29.06±1.57 | 21.13±3.47 | T=-11.36 | <0.001<sup>a</sup> |
| CDR (NC/AD n=31/24) | 0.05±0.15 | 0.92±0.32 | / | / |

Note: Data are presented as the mean ± SD.
NC, normal control; AD, Alzheimer's disease; MMSE, Mini-Mental State Examination; CDR, Clinical Dementia Rating Scale.
<sup>a</sup>The P-value was obtained by the two-sample two-tailed t-test.
<sup>b</sup>The P-value was obtained by the two-tailed Pearson $\chi^2$ test.

**Table 2.** Difference between AD patients and NC on voxel-based ddWD-G. Voxel-based ddWD-G at nine ranges of Euclidean distance were compared between AD patients and NC, adjusted for age, gender, mean frame-wise displacement for head motion. The significance threshold was set at voxel-level p<0.005 (uncorrected) with a cluster-level p<0.05 (FWE-corrected).

| Comparison | Metrics |  | Cluster |  | Peak Voxel |  |  |  |  |
| --- | --- | --- | --- | --- | --- | --- | --- | --- | --- |
| | ddWD | Range (mm) | Size | $P_{FWE}$ | MNI Coordinates | | | T-value | Brain region |
| AD<NC | ddWD-G | 0~20 | 190 | 0.002 | -3 | -15 | 12 | -5.09 | L thalamus |
|  |  |  |  |  | 6 | -18 | 15 | -4.38 | R thalamus |

**Fig 4.**
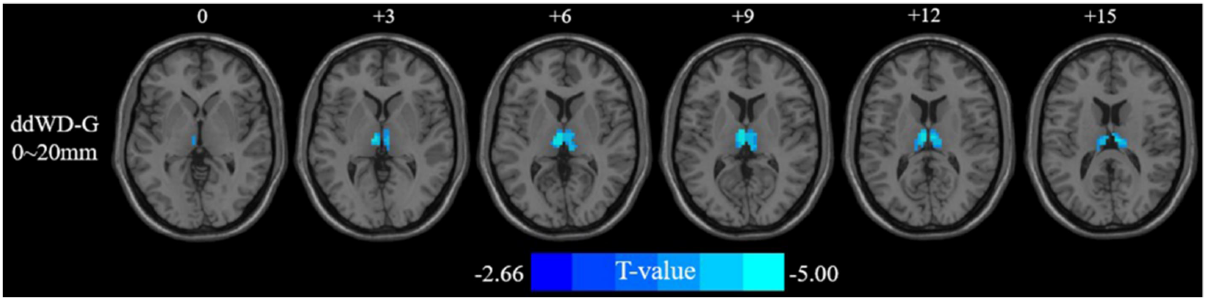
Difference between AD patients and NC on ddWD-G. Voxel-based ddWD-G at nine ranges of Euclidean distance were compared between AD patients and NC, adjusted for age, gender, mean frame-wise displacement for head motion. The significance threshold for ddWD-G set at voxel-level p<0.005 (uncorrected) with a cluster-level p<0.05 (FWE-corrected). Decreased ddWD-G (0∼20 mm) of AD patients were superimposed upon a series of axial slices of standard brain template. The color bar represents the T-value.

### 3.2 Voxel-based ddWD-W

Compared to NC, AD patients showed decreased ddWD-W at 20∼40mm in the left thalamus and subthalamic nucleus. Using this cluster as the seed, we did not find any significant difference between the two groups on the seed-based FCs (Table 3, Fig 5).

**Table 3.** Difference between AD patients and NC on the ddWD-W. Voxel-based ddWD-Ws at eight ranges of Euclidean distance were compared between AD patients and NC, adjusted for age, gender, mean frame-wise displacement for head motion. The significance threshold was set at voxel-level p<0.005 (uncorrected) with a cluster-level p<0.05 (FWE-corrected).

| Comparison | Metrics |  | Cluster |  | Peak Voxel |  |  |  |  |
| --- | --- | --- | --- | --- | --- | --- | --- | --- | --- |
| | ddWD | Range (mm) | Size | $P_{FWE}$ | MNI Coordinates | | | T-value | Brain region |
| AD<NC | ddWD-W | 20~40 | 64 | 0.029 | -17 | -15 | 0 | -2.83 | L thalamus |
|  |  |  |  |  | -11 | -15 | -6 | -3.59 | L subthalamic nucleus |

**Fig 5.**
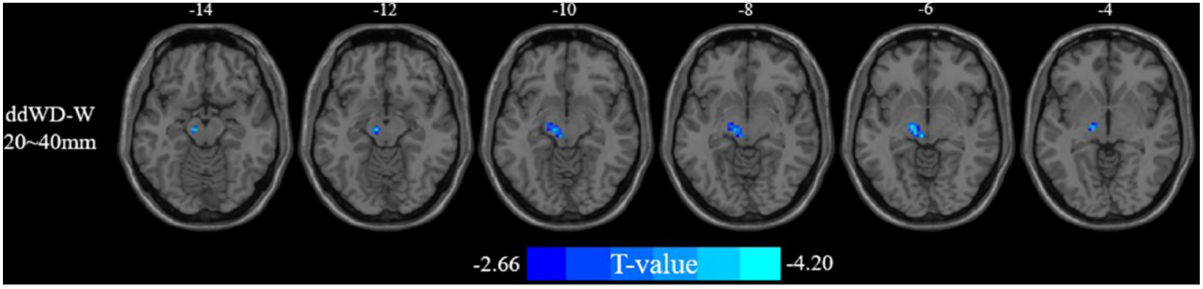
Difference between AD patients and NC on ddWD-W. Voxel-based ddWD-Ws at eight ranges of Euclidean distance were compared between AD patients and NC, adjusted for age, gender, mean frame-wise displacement for head motion. The significance threshold was set at voxel-level p<0.005 (uncorrected) with a cluster-level p<0.05 (FWE-corrected). Decreased ddWD-W (20∼40 mm) of AD patients were superimposed upon a series of axial slices of standard brain template. The color bar represents the T-value.

### 3.3 Voxel-based ddWD-GW and seed-based FC

Compared to NC, AD patients showed increased ddWD-GW at 20∼40mm in the left middle temporal gyrus and middle occipital gyrus. Using this cluster as the seed, we found that the seed-based FCs with the left posterior thalamic radiation were significantly increased in AD patients than NC (Table 4, Fig 6).

**Table 4.**
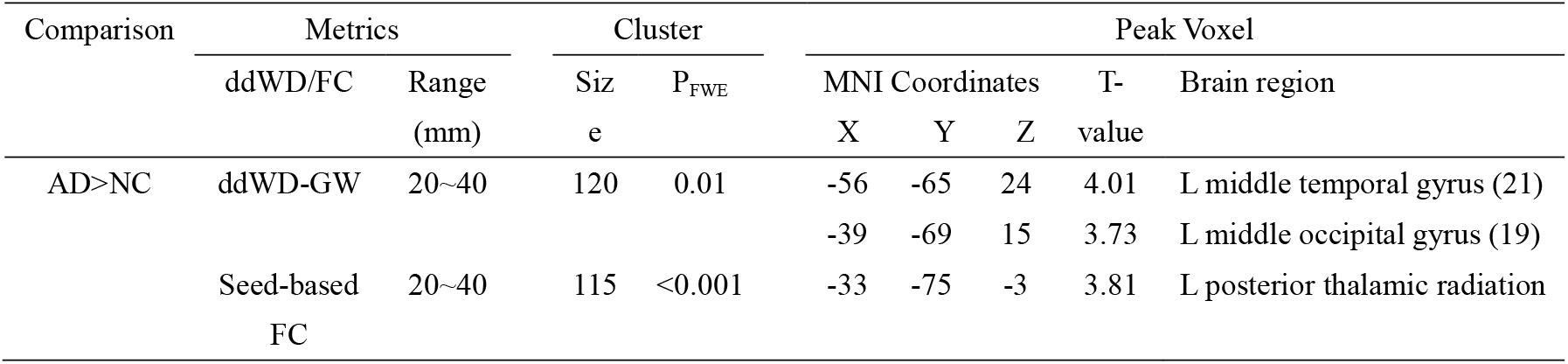
Differences between AD patients and NC on ddWD-GW and the seed-based FC. Voxel-based ddWD-GWs at nine ranges of Euclidean distance were compared between AD patients and NC. Using the resultant supra-threshold cluster as the seed, the seed-based FCs were compared between AD patients and NC. The adjusted covariates included age, gender, mean frame-wise displacement for head motion. The significance threshold was set at voxel-level p<0.005 (uncorrected) with a cluster-level p<0.05 (FWE-corrected).

**Fig 6.**
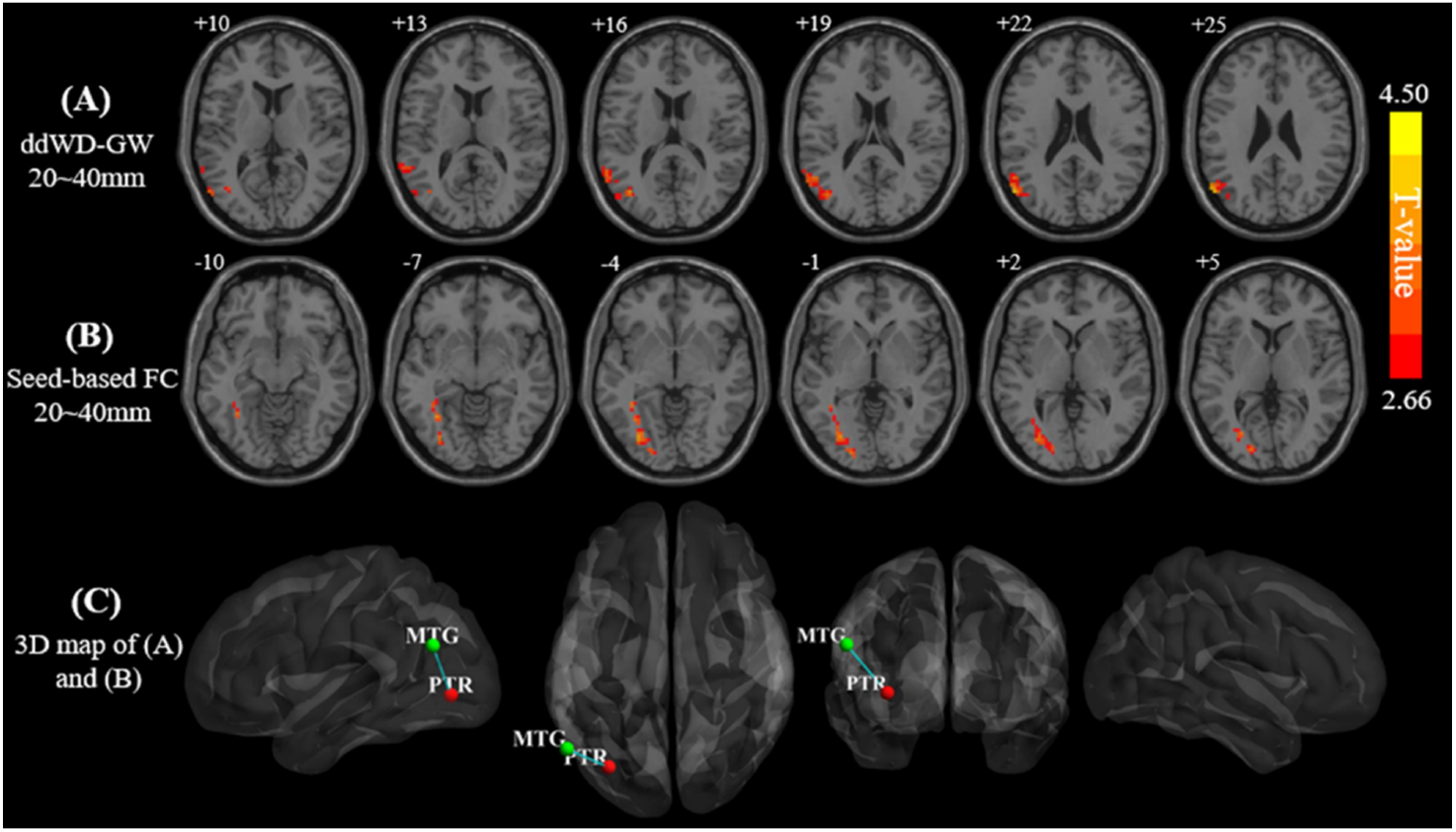
Difference between AD patients and NC on ddWD-GW and the seed-based FC. Voxel-based ddWD-GWs at nine ranges of Euclidean distance were compared between AD patients and NC. Using the resultant supra-threshold cluster as the seed, the seed-based functional connectivity (FC) was compared between AD patients and NC. The adjusted covariates included age, gender, mean frame-wise displacement for head motion. The significance threshold was set at voxel-level p<0.005 (uncorrected) with a cluster-level p<0.05 (FWE-corrected). Increased ddWD-GW at 20∼40 mm A) and increased seed-based FCs B) of AD patients were superimposed upon a series of axial slices of standard brain template. The color bar represents the T-value. The spatial distribution of the seed (in red) and seed-based FCs (in green) was shown in C), with a sphere representing a cluster, centering on the cluster’s peak voxel. The line between the seed and the cluster indicated their FC, and the thickness of the line represented the T-value of group comparison on FC. Abbreviation: MTG middle temporal gyrus, PTR posterior thalamic radiation.

### 3.4 Voxel-based ddWD-WG and seed-based FC

Compared to NC, AD patients showed decreased ddWD-WGs at two ranges (40∼60mm and 60∼80mm) in the left superior corona radiation, and increased ddWD-WGs at four ranges (20∼40mm, 40∼60mm, 60∼80mm, and 80∼100mm) in the left posterior thalamic radiation and right sagittal stratum (Table 5, Fig 6). Using the supra-threshold clusters as the seeds, we found that the FCs of two clusters were significantly increased in AD patients than NC. The two clusters were both located in the right sagittal stratum but differed on the distance ranges. The seed-based FCs of the cluster at 60∼80mm were located in the bilateral supplementary motor area, left middle cingulate gyrus, right superior frontal gyrus, and right caudate. The seed-based FCs of the cluster at 80∼100mm were located in the bilateral supplementary motor area, left insula, left anterior cingulate cortex, right medial superior frontal gyrus, right superior frontal gyrus (Table 5, Fig 7).

**Table 5.** Difference between AD patients and NC on ddWD-WG and the seed-based FC. Voxel-based ddWD-WGs at nine ranges of Euclidean distance were compared between AD patients and NC. Using the resultant supra-threshold clusters as the seed, the seed-based functional connectivity (FC) was compared between AD patients and NC. The adjusted covariates included age, gender, mean frame-wise displacement for head motion. The significance threshold was set at voxel-level p<0.005 (uncorrected) with a cluster-level p<0.05 (FWE-corrected).

| Comparison | Metrics |  | Cluster |  | Peak Voxel |  |  |  |  |
| --- | --- | --- | --- | --- | --- | --- | --- | --- | --- |
|  | ddWD/FC | Range<br>(mm) | Size | P<sub>FWE</sub> | MNI Coordinates |  |  | T-value | Brain region (BA) |
|  |  |  |  |  | X | Y | Z |  |  |
| AD<NC | ddWD-WG | 40~60 | 55 | 0.034 | -33 | -12 | 21 | -4.13 | L superior corona radiation |
|  | ddWD-WG | 60~80 | 59 | 0.022 | -30 | -3 | 30 | -3.68 | L superior corona radiation |
| AD>NC | ddWD-WG | 20~40 | 64 | 0.022 | -39 | -57 | -3 | 3.73 | L posterior thalamic radiation |
|  | ddWD-WG | 40~60 | 75 | 0.008 | -39 | -57 | -3 | 3.66 | L posterior thalamic radiation |
|  | ddWD-WG | 60~80 | 49 | 0.047 | -36 | -63 | 9 | 3.94 | L posterior thalamic radiation |
|  |  |  | 45 (Seed1) | 0.065 | 36 | -39 | 0 | 3.83 | R sagittal stratum |
|  | ddWD-WG | 80~100 | 55 (Seed2) | 0.029 | 36 | -39 | 3 | 3.87 | R sagittal stratum |
| AD>NC | Seed1-based FC | 60~80 | 79 | 0.014 | 15 | 21 | -6 | 3.91 | R caudate |
|  |  |  | 211 | <0.001 | -6 | -6 | 60 | 4.46 | L supplementary motor area (6) |
|  |  |  |  |  | -6 | 6 | 36 | 4.08 | L middle cingulate cortex (24) |
|  |  |  | 67 | 0.03 | 21 | 3 | 57 | 3.93 | R superior frontal gyrus (8) |
|  |  |  |  |  | 12 | -15 | 69 | 3.49 | R supplementary motor area (6) |
|  | Seed2-based FC | 80~100 | 64 | 0.038 | -39 | -3 | 9 | 3.61 | L insula |
|  |  |  | 569 | <0.001 | 21 | 42 | 36 | 4.60 | R superior frontal gyrus (9) |
|  |  |  |  |  | 12 | 30 | 57 | 4.37 | R superior medial frontal gyrus (8) |
|  |  |  |  |  | -1 | 39 | 24 | 4.34 | L anterior cingulate cortex (32) |
|  |  |  | 76 | 0.018 | -6 | -6 | 60 | 4.45 | L supplementary motor area (6) |
|  |  |  |  |  | 12 | -15 | 69 | 3.51 | R supplementary motor area (6) |
|  |  |  | 64 | 0.038 | 27 | 12 | 57 | 3.83 | R superior frontal gyrus (8) |

**Fig 7.**
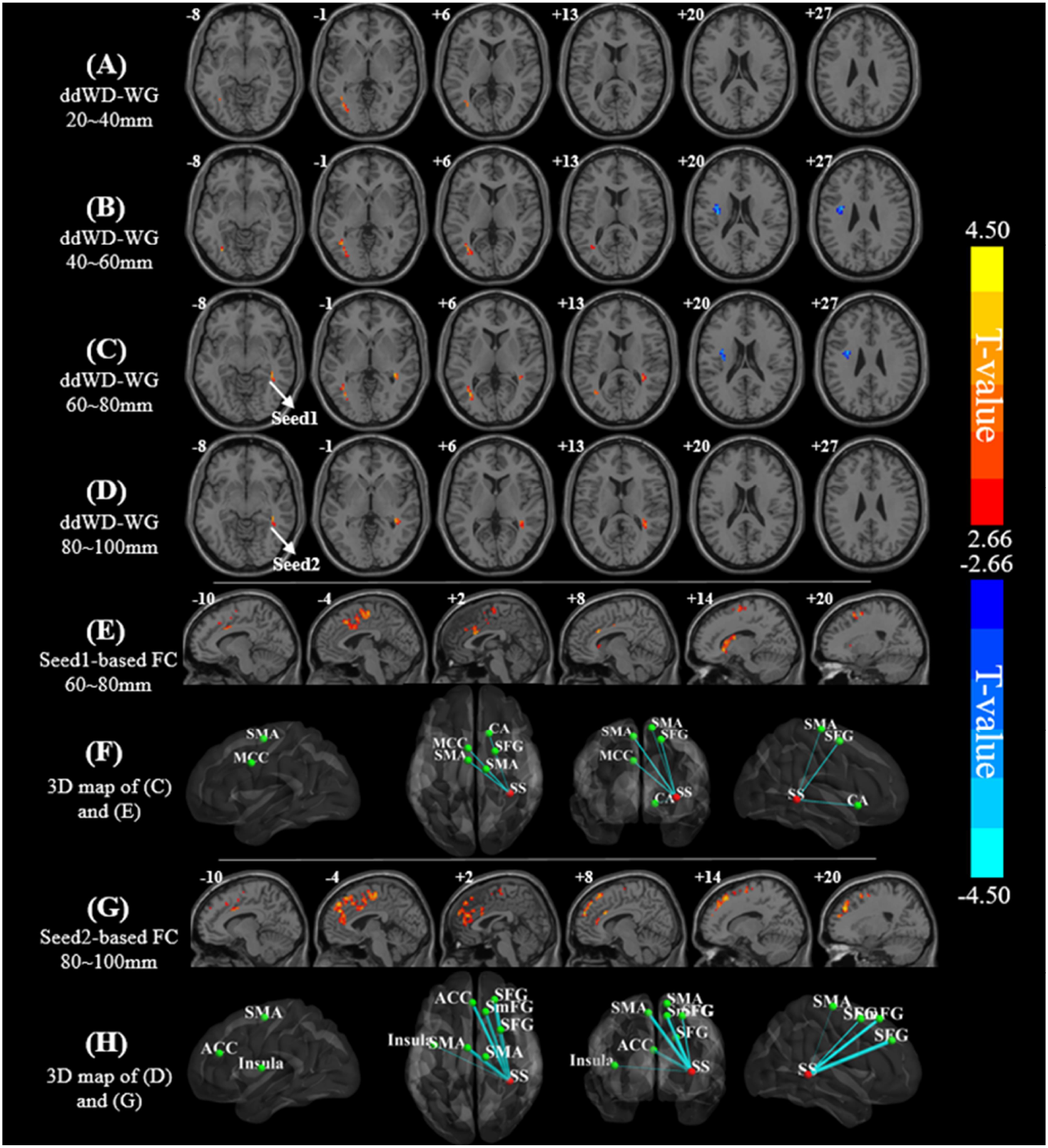
Difference between AD patients and NC on ddWD-WG and seed-based FC. Voxel-based ddWD-WGs at nine ranges of Euclidean distance were compared between AD patients and NC. Using the resultant suprathreshold cluster as the seeds, the seed-based functional connectivity (FC) was compared between AD patients and NC. The adjusted covariates included age, gender, mean frame-wise displacement for head motion. The significance threshold was set at voxel-level p<0.005 (uncorrected) with a cluster-level p<0.05 (FWE-corrected). Increased ddWD-WG metrics (in red) of AD patients were found at A) 20∼40 mm, B) 40∼60mm, C) 60∼80mm, and D) 80∼100mm, while decreased ddWD-WD metrics (in blue) of AD patients were found at B) 40∼60mm and C) 60∼80mm. Using the two clusters in C) and D) as the seeds, we found significantly larger seed-based FCs in AD than NC, with the seed1-based FCs shown in E) and seed2-based FCs shown in G. The color bar represents the T-value (positive T-value in red, negative T-value in blue). The spatial distribution maps of two seeds (in red) and seed-based FCs (in green) were shown in F) and H), respectively, with each sphere representing a cluster, centering on the cluster’s peak voxel. The line between the seeds and the clusters indicated their FC, and the thickness of the line represented the T-value of group comparison on FC. Abbreviation: CA caudate nucleus, SMA supplementary motor area, MCC middle cingulate gyrus, SFG superior frontal gyrus, SS sagittal stratum, ACC anterior cingulate gyrus, SmFG superior medial frontal gyrus.

## 4. Discussion

### 4.1 Decreased ddWD-G

Our additional analysis of voxel-based GM volumes demonstrated significantly smaller GM volumes in the hippocampus, middle and inferior temporal gyrus in AD than NC, consistent with the classical locations of GM atrophy of AD (Basso et al., 2006; Nho et al., 2012). Our results showed that AD patients had decreased ddWD-G at 0∼20mm in the bilateral thalamus than NC. The thalamus, along with the hippocampus and posterior cingulate cortex, were involved in the memory system (Aggleton, Pralus, Nelson, & Hornberger, 2016); the atrophy of bilateral thalamus was also observed in AD patients (Agosta et al., 2012). Dai and colleagues showed that the FCs between the thalamus and the neighboring regions at a short distance of 30∼40 mm were significantly smaller in AD patients than NC (Z. J. Dai et al., 2015), consistent with our finding.

### 4.2 Decreased ddWD-W

We found that AD patients had decreased ddWD-W at 20∼40mm in the thalamus and subthalamic nucleus, compared to NC. A previous study showed decreased fractional anisotropy values and smaller volume in the thalamus in AD patients (Stepan-Buksakowska et al., 2014; Zarei et al., 2010). Neuropathological changes of neurofibrillary tangles and deposition of amyloid have been found in the thalamus and subthalamic nucleus of AD patients (Braak & Braak, 1991; Mattila, Togo, & Dickson, 2002). As the BOLD signal was dependent on the structural integrity, the decreased ddWD-W in the thalamus and subthalamic nucleus observed in our study might reflect the neuropathological damage in the two areas.

### 4.3 Increased ddWD-GW and seed-based FC

Several theories proposed a compensation mechanism in the aging brain, suggesting that the plastic brain could reorganize functional circuits to offset the age-related neural inefficiency (Park & Reuter-Lorenz, 2009; Reuter-Lorenz & Cappell, 2008). Converging evidence showed that compared to younger adults, older adults showed enhanced neural activity during performing various tasks (Cabeza, Anderson, Locantore, & McIntosh, 2002; Reuter-Lorenz et al., 2000), and increased FCs in the various cortical areas (Ferreira & Busatto, 2013; Tomasi & Volkow, 2012). Likewise, the compensation mechanism possibly existed in AD patients. Previous studies showed that the neural activities of AD patients were stronger than NC in the dorsolateral prefrontal cortex, insula, supplementary motor areas during various tasks (Gould et al., 2006; Z. Q. Wang et al., 2011; Yetkin, Rosenberg, Weiner, Purdy, & Cullum, 2006). The increased FCs in the salience and executive networks were also found in AD patients, which possibly compensated for cognitive efficiency decline (Y. F. Zhang et al., 2016). However, no prior study had used the WM network metrics to demonstrate the compensation phenomenon in AD patients.

Our study showed that AD patients had larger ddWD-GW at 20∼40mm in the left middle temporal and occipital cortices than NC. Using this cluster as the seed, we found that the FCs between this seed and the left posterior thalamic radiation were larger in AD patients than NC. The middle temporal and occipital cortices were involved in visuospatial processing, and the posterior thalamic radiation connected the thalamus with the visual cortices (Prvulovic et al., 2002a; Tuch et al., 2005; Vannini et al., 2008). Prvulovic and colleagues found that the activation of the left occipito-temporal cortex was significantly higher in AD patients than NC, possibly indicating an attempt to offset the neural inefficiency (Prvulovic et al., 2002b). Our results of increased ddWD and FCs in these areas might reflect the compensation mechanism of AD patients in the visuospatial function.

### 4.4 Decreased and increased ddWD-WG and seed-based FC

#### Decrease ddWD-WG metrics in AD than NC

The present study showed decreased ddWD-WG at 40∼60 mm and 60∼80mm in the left superior corona radiation in AD patients than NC. Leunissen and colleagues performed a tractography study that showed the superior corona radiation connected the basal ganglia and thalamus to the frontal areas (Leunissen et al., 2014). The structural integrity of superior corona radiation was correlated with processing speed and flexibility across age (Leunissen et al., 2014; Moeller, Willmes, & Klein, 2015; Radoeva et al., 2020; Sasson, Doniger, Pasternak, Tarrasch, & Assaf, 2012). In AD, the structural integrity of superior corona radiation was disrupted, evidenced with decreased fractional anisotropy (Birdsill et al., 2014; Ouyang et al., 2015; Yin et al., 2015). Our results regarding the decreased ddWD-WGs in the left superior corona radiation possibly reflected the damage in this fiber bundle and deficits in relevant cognitive functions in AD patients.

#### Increased ddWD-WG metrics in AD than NC

Compared to NC, AD patients showed increased ddWD-WG at 20∼40mm, 40∼60mm, and 60∼80mm in the left posterior thalamic radiation. This location was consistent with our prior finding of increased seed-based FCs of AD in the left posterior thalamic radiation, with the seed located in the left middle temporal-occipital cortices. As mentioned above, the posterior thalamic radiation is involved in visuospatial processing by connecting the thalamus and the visual cortices (Tuch et al., 2005). The increased ddWD-WG in the left posterior thalamic radiation possibly reflected the compensation of visuospatial functional circuits in AD patients.

We also found increased ddWD-WG at 60∼80mm and 80∼100mm in the right sagittal stratum in AD patients than NC. The sagittal stratum contains the middle and inferior longitudinal fascicles and the inferior fronto-occipital fascicle, as well as the optic radiation (Di Carlo et al., 2019). By connecting with the extensive areas in the frontal, temporal, parietal, and occipital areas, the sagittal stratum was involved in attention (Altieri et al., 2019), executive control, and affective process (Chan-Seng, Moritz-Gasser, & Duffau, 2014; Ebeling & Reulen, 1988; Mandonnet et al., 2017; Rayhan et al., 2013).

Using the clusters in the sagittal stratum as the seeds, we found increased seed-based FCs in the regions of two networks, i.e., the salience network (insula and anterior cingulate cortex), and the executive network (medial frontal and superior frontal gyri) for AD patients. Several studies on the GM functional network changes in AD patients demonstrated increased FCs in the salience and executive networks (Agosta et al., 2012; K. Wang et al., 2007; Zhou et al., 2010). The salience network processed and integrated internal and external inputs to facilitate cognitive control, which was the undertaking of executive network (Seeley et al., 2007; Sridharan, Levitin, & Menon, 2008). The neural activities in both networks were in anti-correlation with default mode network whose degeneration was a hallmark of AD pathology (Fox et al., 2005; Greicius, Srivastava, Reiss, & Menon, 2004). A compensatory theory was proposed to explain such dichotomy, suggesting that accompanied by the inactive task-negative system, the task-positive systems became more active to offset the inefficient neural processes, albeit possibly inadequate eventually (Agosta et al., 2012; Brier et al., 2012; Dennis & Thompson, 2014). Our finding provided new evidence for the compensatory mechanism of AD, and identified the sagittal stratum as a key WM structure involved in it.

### 4.5 Useful biomarker of ddWD metrics

In the present study, the analysis with voxel-based WD metrics did not show any significant results, while the analysis with voxel-based ddWD-GW and ddWD-WG identified the compensatory networks in AD. A recent study suggested an increased coupling between functional and structural connectivity networks in AD (Z. Dai et al., 2019). The FCs between two tissues were more effective than the FCs within GM to capture the pathological characteristics of AD (Zhao et al., 2019). Moreover, we noted that the changes of ddWD were relying on the metric type, location, and Euclidean distance. In our study, the decreased metrics were mainly short ranged (0∼40mm) ddWD-G/W in the thalamus; meanwhile the increased metrics were short-to-medium ranged (20∼100mm) ddWD-GW/WG between certain WM tracts and GM areas. Dai and colleagues showed smaller short-ranged FC strength (30-40 mm) in the thalamus and prominent long-ranged FCs disruption in AD (Z. J. Dai et al., 2015). However, the short- and medium-ranged FCs were enhanced in the participants with subjective cognitive decline and AD, suggesting a compromise of the compensatory neural process on the benefit and cost (H. Chen et al., 2020; Liu et al., 2014).

## 5. Limitation

The present study had a few limitations. Firstly, the number of eligible participants of our study was limited, as 30 AD patients and 37 NC in the ADNI database met our inclusion criteria. Secondly, given that the complete neuropsychological information was not available for each participant from the ADNI database, we could not perform correlation analyses to examine the cognitive relationships of functional network metrics. Finally, the ddWD metrics that we employed were based on static FC. However, the temporal synchronization between WM and GM was dynamic, and the causal relationship between the two tissues would be of much interest for future exploration in AD patients.

## 6. Conclusion

Compared to NC, AD patients showed both decreased and increased ddWD metrics. The distinct change patterns of ddWD might indicate the co-existence of neurodegnerative and compensatory mechanisms in AD. The change pattern of ddWD was dependent on the metric type, location, and Euclidean distance, as the increased were the short-to-medium ranged ddWD metrics between WM tracts and GM areas, possibly reflecting a compromise between benefit and cost.

## 7. Author contributions

Xingxing Zhang: conceptualization, methodology, writing the original draft. Qing Guan: review & editing. Debo Dong: review & editing. Fuyong Chen: review & editing. Jing Yi: review & editing. Yuejia Luo: review & editing. Haobo Zhang: conceptualization, methodology, writing, review & editing.

## 8. Acknowledgements

This study was supported by the National Natural Science Foundation of China (No. 31700960, 32071100), the Natural Science Foundation of Guangdong Province of China (2020A1515011394), the Shenzhen Fundamental Research General Project (JCYJ20190808121415365), the Shenzhen-Hong Kong Institute of Brain Science-Shenzhen Fundamental Research Institutions grant (2019SHIBS0003), and the National Key Research and Development Program of China (2018YFC1315205). The funders had no role in study design, data collection and analysis, decision to publish, or preparation of the manuscript.

The data used in this study was from the Alzheimer’s Disease Neuroimaging Initiative (ADNI) (National Institutes of Health Grant U01 AG024904) and DOD ADNI (Department of Defense award number W81XWH-12-2-0012). ADNI is funded by the National Institute on Aging, the National Institute of Biomedical Imaging and Bioengineering, and through generous contributions from the following: AbbVie, Alzheimer’s Association; Alzheimer’s Drug Discovery Foundation; Araclon Biotech; BioClinica, Inc.; Biogen; Bristol-Myers Squibb Company; CereSpir, Inc.; Cogstate; Eisai Inc.; Elan Pharmaceuticals, Inc.; Eli Lilly and Company; EuroImmun; F. Hoffmann-La Roche Ltd. and its affiliated company Genentech, Inc.; Fujirebio; GE Healthcare; IXICO Ltd.; Janssen Alzheimer Immunotherapy Research & Development, LLC.; Johnson & Johnson Pharmaceutical Research & Development LLC.; Lumosity; Lundbeck; Merck & Co., Inc.; Meso Scale Diagnostics, LLC.; NeuroRx Research; Neurotrack Technologies; Novartis Pharmaceuticals Corporation; Pfizer Inc.; Piramal Imaging; Servier; Takeda Pharmaceutical Company; and Transition Therapeutics. The Canadian Institutes of Health Research is providing funds to support ADNI clinical sites in Canada. Private sector contributions are facilitated by the Foundation for the National Institutes of Health (www.fnih.org). The grantee organization is the Northern California Institute for Research and Education, and the study is coordinated by the Alzheimer’s Therapeutic Research Institute at the University of Southern California. ADNI data are disseminated by the Laboratory for Neuro Imaging at the University of Southern California.

## 10. Data for reference

All the data was obtained from openly available data sets. Code will be deposited in an open access platform, upon acceptance of this manuscript.

