## Supplemental Fig s1, Supplemental Fig s2, and Supplemental Table s1 for "Compensatory reconfiguration of functional networks between white matter and grey matter in Alzheimer’s disease"

Supplementary materials

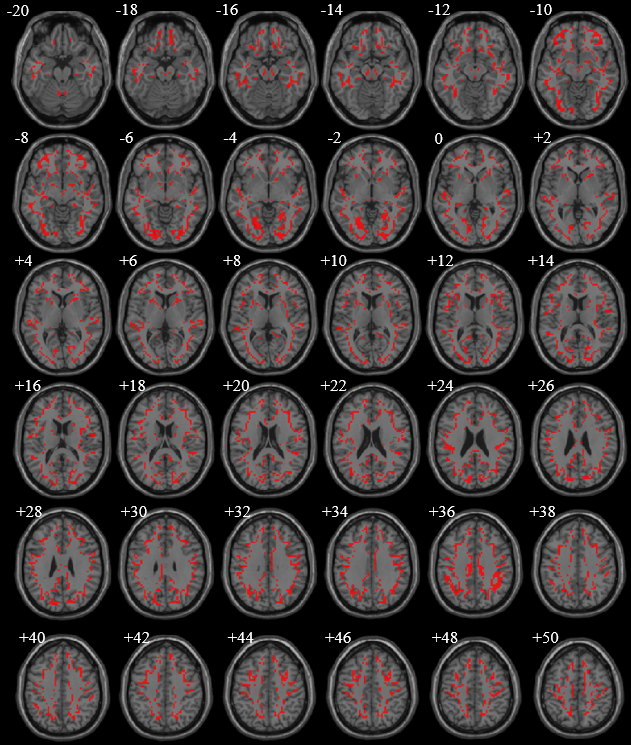

**Fig s1. The overlapped voxels between gray-matter mask and white-matter mask.** These voxels (red) were located on the boundary of the GM and WM masks, with a GM probability >0.2 and WM probability >0.6. They were superimposed upon a series of axial slices of standard brain template. Considering that the attribution of these voxels is still controversial, we did not include them in the GM and WM masks for subsequent analysis.

**Measured the changes of GM and WM volume in AD patients**

Image acquisition

All participants were scanned with a 3.0 T Philips scanner (Jack et al., 2008). The T1-weighted images were acquired using the following parameters: TR = 6700 ms, TE = 3.1 ms, FA = 9°, slice thickness = 1.2 mm, matrix = 256 × 256 × 170. The rs-fMRI images were obtained using an echo-planar imaging (EPI) sequence with the following parameters:

repetition time (TR) = 3000 ms, echo time (TE) = 30 ms, flip angle = 80°, number of slices = 48, slice thickness = 3.313 mm, voxel size = 3 mm × 3 mm × 3 mm, voxel matrix = 64 × 64, and total volume = 140.

T1 Image processing

We applied the voxel-based morphometry approach in image processing by running the Statistical Parametric Mapping software (SPM12, *http://*[*www.fil.ion.ucl.ac.uk/spm*](http://www.fil.ion.ucl.ac.uk/spm)). After the visual inspection for structural abnormalities, the T1-weighted images were segmented using the hidden Markov random field. A series of customized templates and flow fields were then generated using the diffeomorphic anatomical registration through an exponentiated lie algebra (DARTEL) registration method. The images were then registered to the customized templates and then normalized to the Montreal Neurological Institute (MNI) space. Finally, the images were smoothed with a 12-mm full-width at half-maximum Gaussian kernel.

Voxel-based GM volume

Compared with normal controls, AD patients showed significantly smaller GM volumes in the bilateral hippocampi, middle temporal gyri, and amygdala.

| Comparison | Metrics | |  | Cluster | |  | Peak Voxel | | | | |
| --- | --- | --- | --- | --- | --- | --- | --- | --- | --- | --- | --- |
|  |  |  |  | Size | P^FWE^ |  | MNI Coordinates  X Y Z | | | T-  value | Brain region (BA) |
| AD<NC | GM volume | |  | 13081 | 0.001 |  | -54 | -51 | 2 | -6.70 | L middle temporal gyrus (21) |
|  |  |  |  |  |  |  | -26 | -3 | -17 | -6.38 | L amygdala |
|  |  |  |  |  |  |  | -27 | -32 | -5 | -5.73 | L hippocampus |
|  |  |  |  |  | 0.07 |  | 20 | -3 | -15 | 4.81 | R amygdala |
|  |  |  |  |  |  |  | 32 | -27 | -6 | -4.44 | R hippocampus |

**Table s1. Difference between AD patients and normal controls on voxel-based GM volume.** Voxel-based GM volume were compared between AD patients and normal controls, adjusted for age, gender, and total intracranial volume. The significance threshold for GM volume set at voxel-level p<0.005 (uncorrected) with a cluster-level p<0.05 (FWE-corrected).

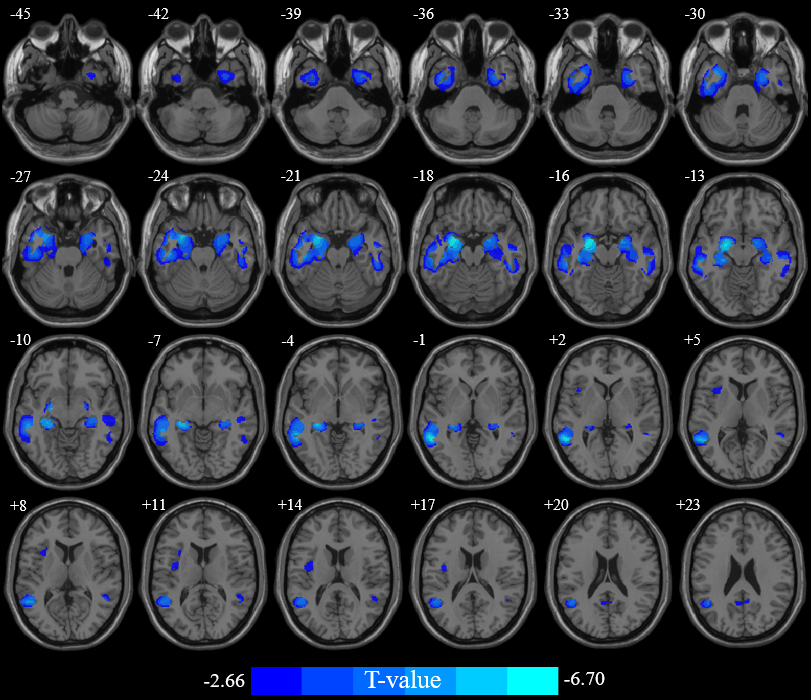

**Fig s2. Difference between AD patients and normal controls on voxel-based GM volume (GMV).** Voxel-based GM volume were compared between AD patients and normal controls, adjusted for age, gender, and TIV. The significance threshold for GM volume set at voxel-level p<0.005 (uncorrected) with a cluster-level p<0.05 (FWE-corrected). The significant results for GM volume were superimposed upon a series of axial slices of standard brain template. The color bar represents the T-value.
